# Rapid Structure-Function Insights via Hairpin-Centric Analysis of Big RNA Structure Probing Datasets

**DOI:** 10.1101/2021.04.27.441661

**Authors:** Pierce Radecki, Rahul Uppuluri, Sharon Aviran

**Affiliations:** Biomedical Engineering Department and Genome Center, University of California at Davis, Davis, CA, 95616, USA

## Abstract

The functions of RNA are often tied to its structure, hence analyzing structure is of significant interest when studying cellular processes. Recently, large-scale structure probing (SP) studies have enabled assessment of global structure-function relationships via standard data summarizations or local folding. Here, we approach structure quantification from a hairpin-centric perspective where putative hairpins are identified in SP datasets and used as a means to capture local structural effects. This has the advantage of rapid processing of big (e.g., transcriptome-wide) data as RNA folding is circumvented, yet it captures more information than simple data summarizations. We reformulate a statistical learning algorithm we previously developed to significantly improve precision of hairpin detection, then introduce a novel nucleotide-wise measure, termed the hairpin-derived structure level (HDSL), which captures local structuredness by accounting for the presence of likely hairpin elements. Applying HDSL to data from recent studies recapitulates, strengthens, and expands on their findings which were obtained by more comprehensive folding algorithms, yet our analyses are orders of magnitude faster. These results demonstrate that hairpin detection is a promising avenue for global and rapid structure-function analysis, furthering our understanding of RNA biology and the principal features which drive biological insights from SP data.

## INTRODUCTION

RNA structure is driven primarily by the complementarity of nucleotide bases comprising it, which allows for hydrogen bonding between various segments of the molecule. Intramolecular base pairing, combined with the flexible and single-stranded nature of the molecule’s backbone, allows for intricate secondary and tertiary structural elements. These structures, as well as their ability to dynamically change between relevant configurations, are known to play central roles in almost every facet of cellular regulation (1–6). Understanding the structures of RNA is therefore important, which has led to an explosion of methods which probe (7–17), computationally predict (18–28), and interpret them in various contexts (1, 5, 29–35).

Structure probing (SP) experiments currently provide the most practical approach for measuring RNA structures in their natural environment. These experiments work by exposing RNA to chemicals, enzymes, or photons which react differentially with parts of the molecule depending on their structural context (for example, paired/unpaired nucleotides or ds/ssRNA) (7, 8, 10–13, 36, 37). Specific protocols vary, but typically the probing reaction induces changes to the RNA bases or backbone which are detected via sequencing or electrophoresis as mutations or truncations (38, 39). The rate of mutation or truncation at a particular nucleotide is used to summarize that nucleotide’s reactivity with the probe (40). These data contain critical information on the structural conformation of an RNA, and incorporating them as soft constraints within thermodynamics-based folding algorithms greatly improves their accuracy (18, 26, 41).

Next-generation sequencing has allowed SP experiments to scale to the level of the whole cell (i.e., transcriptome-wide). Exploration of these data have typically begun with straightforward global-level quantifications and simple comparisons (11, 42–46). More recent studies expanded the intricacy of structural analysis to disentangle the dynamic functional roles of RNA structure in fundamental cellular processes (47). For example, Saha et al. compared reactivity profiles in the vicinity of spliced introns and retained introns, and found evidence of increased structure upstream and decreased structure downstream of retained introns (48). Yang et al. characterized structural impacts on miRNA-mediated mRNA cleaving by computing mean reactivity and mean base-pairing probability profiles around miRNA target sites, which illuminated a strong connection between transcript cleavage and unpaired bases immediately downstream of the miRNA target site (49). Work by Mustoe et al. (30) and Mauger et al. (50) have linked changes in gene expression within *E. coli* and human cells to the structural dynamics within coding sequences and UTRs as quantified by local median reactivities. Twittenhoff et al. (51) performed structure probing of *Y. pseudotuberculosis* at different temperatures and used averaged reactivity scores to highlight differential structure changes due to temperature in 5’UTRs versus coding regions in addition to using condition-wise reactivity differences to identify temperature-sensitive genes.

A common theme to such studies is the quantification of local “structuredness” and comparisons of it at global scales. To this end, measures of structure are typically founded on basic statistical summarization of reactivities, sometimes combined with data-directed thermodynamics-based folding algorithms to quantify base-pairing probabilities. Current state-of-the-art algorithms for predicting base-pairing probabilities (and specific RNA structures) are founded on dynamic programming strategies and a nearest neighbor thermodynamic model (NNTM) (52, 53). Although relatively efficient, these scale as *O*(*L*^3^) with the length of an RNA, meaning that complete folding analyses of long RNA transcripts are often computationally infeasible. NNTM-based processing (i.e., RNA folding and computation of base-pairing probabilities) of the massive data associated with recent studies is thus challenging. As a consequence, transcriptome-wide studies have typically utilized ad-hoc folding strategies which attempt to strike a balance between computational overhead and prediction quality by locally folding pre-screened candidate regions or rolling windows of long transcripts. Even with such compromises, *in silico* analyses can take days to complete, depending on the scale of the experiment. The process itself is also susceptible to high error rates especially in molecules with multiple stable conformations (54). It is worth noting that some of the aforementioned experiments relied solely on simple reactivity summarization; nevertheless, even in such situations, detections are typically limited to the most pronounced effects. More sophisticated analysis which accounts for structure in addition to reactivity has the potential to refine such findings and expand on them (55, 56). This highlights a need for methods capable of rapidly extracting pertinent structural information from reactivity data.

Motivated by this need, we harnessed *patteRNA*, an NNTM-free method we previously introduced for rapidly mining structural motifs (57, 58), to quantify global trends in RNA structure dynamics from SP data. Briefly, the method works in two phases: training and scoring. The training phase learns a hidden Markov model (HMM) of secondary structure and a Gaussian mixture model (GMM) of the reactivity distributions of paired and unpaired nucleotides. The learned distributions are used to score sites for their likelihood to harbor any target structural motif (see **Figure 1A**). *patteRNA* can automatically process data from any type of SP experiment. Although we previously demonstrated that *patteRNA* accurately detects structural motifs in diverse datasets, we found that there was nevertheless room for significant improvement. Namely, there was a need for improved precision of motif detection, particularly pertaining to the vast search space encountered in transcriptome-wide experiments. Additionally, we found that our method, although suitable for comparative analysis of motifs (58), did not provide a clear quantitative framework for making practical and direct structural inferences in large datasets.

**Figure 1.**
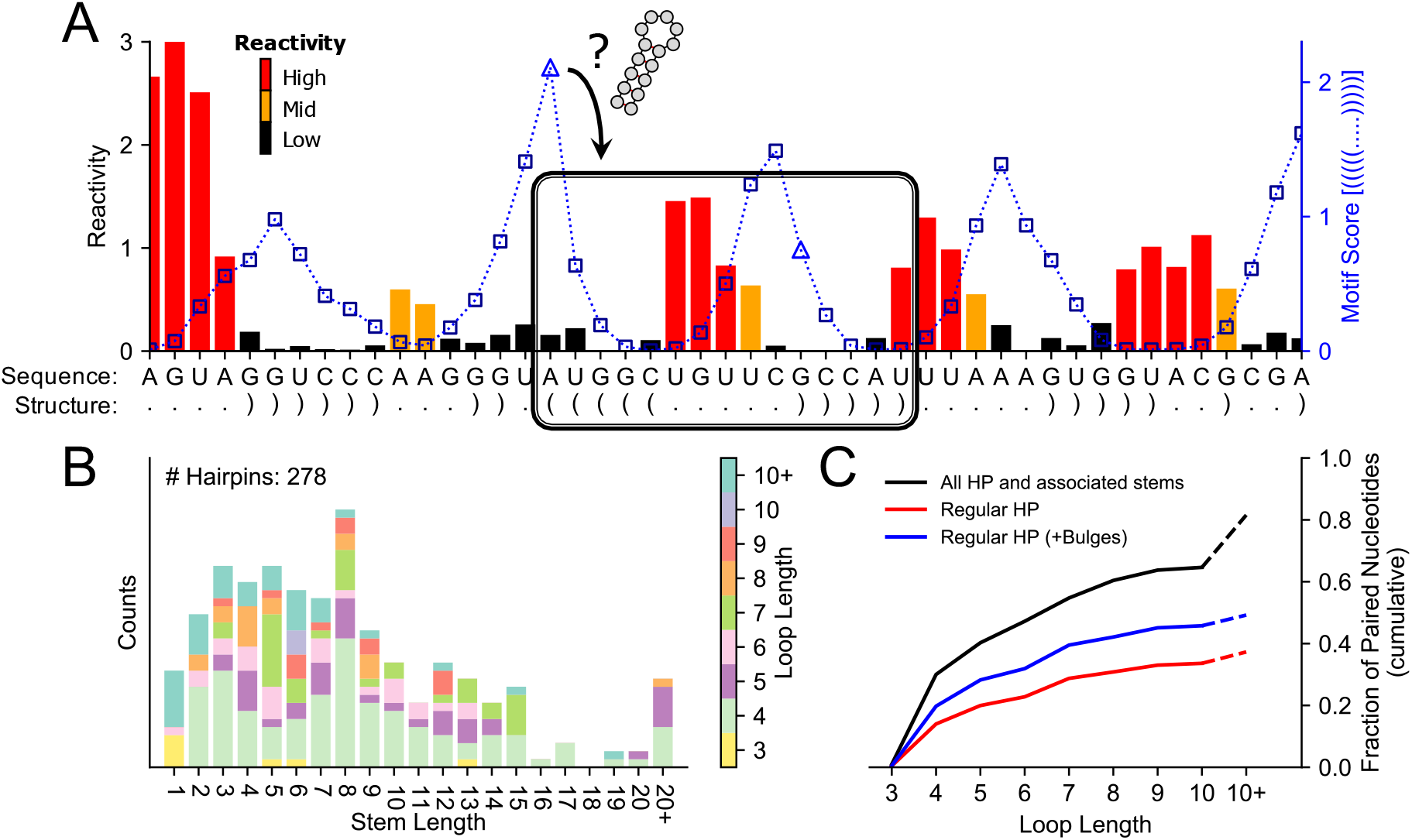
Identification of structural motifs in probing data and representation of hairpins in structures. (A) Schematic illustrating reactivity profile (black, yellow, red) for a region against the corresponding *patteRNA* score profile (blue) when mining for a hairpin with loop length 5 and stem length 5 (dot-bracket: “(((((.....)))))”). The score profile represents the likelihood of the target motif occurring at the site obtained by using the current nucleotide as the start (left side) of a sliding window. This profile achieves a maximum at the true positive site of the hairpin (indicated as black box). Locations which do not satisfy sequence constraints necessary for the base pairs of the motif are denoted by square-shaped markers on the score profile, and vice versa for triangle-shaped markers. Data shown are SHAPE-Seq reactivities from the 23S rRNA of *E. coli* (nt 2531-2576) (41). Reactivities are color coded according to their magnitude (high: > 0.7; mid: > 0.3 and ≤ 0.7; low: ≤ 0.3). (B) Distribution of hairpin stem and loop lengths in a diverse set of structured RNAs (referred to as the Weeks set – see Methods). The vast majority of hairpins have stem lengths shorter than 15 nt and loop lengths between 3 and 10 nt. (C) Fraction of paired nucleotides in the Weeks set which can be represented as belonging to a regular hairpin (red), a regular hairpin with up to one or two bulges of length 1–5 nt (blue), or any/all type of hairpin and associated stems (black).

In this article, we expand and improve the capabilities of *patteRNA* and demonstrate that motif detection can be used to rapidly quantify RNA structuredness in SP datasets. As a first step, we investigate the properties of hairpin elements in RNA structures and their prevalence among all structural elements. We then present an improved unsupervised training approach which yields more accurate motif detection, especially for hairpins, and benchmark it against diverse types of data. Next, we describe a novel measure, the hairpin-derived structure level (HDSL), which uses *patteRNA*’s detected hairpins to quantify the local structure context around nucleotides. We apply HDSL to three recent large-scale SP datasets to demonstrate that our hairpin-driven analysis is 1) capable of recapitulating, strengthening, and expanding on previously detected structural effects and 2) orders of magnitude faster than comparable NNTM-based routines. Simply put, our method bridges the gap between quick but naïve data summarization and intensive but more sophisticated folding-based analysis to provide rapid structure-aware interpretations. Overall, the results of our work also serve to further our understanding of the ways in which diverse SP datasets can be automatically quantified and interpreted without dependence on the assumptions driving NNTM predictions and the complexities associated with them.

## MATERIAL AND METHODS

### Hairpin Counting and Quantification in Known Structures

#### All hairpins (hairpins and associated stems, with or without bulges)

Hairpins in reference dot-bracket structures were retrieved by first identifying hairpin-loops, and then backtracking to determine the full stem length. Hairpin loops are defined as locations in the dot-bracket structures where a base pair flanks a sequence of unpaired states of any length (for example, “(…)” or “(......)”). Once a hairpin loop is identified, the stem length is determined by walking along the structure in both directions until a branching base pair is detected (i.e., a “)” to the left of the stem-loop or a “(“ to the right). At this point, the stem length is called as the number of nested base pairs before the first branching base pair on either side of the stem. As a consequence, bulges and internal loops are generally ignored, so long as they occur before a branching base pair. Loops which are involved in pseudo-knotted base-pairing are treated as unpaired loops for the purpose of hairpin identification.

#### Regular hairpins (hairpins without bulges)

We defined regular hairpins as hairpins having a stem length between 4–15 nt and loop length between 3–10 nt with no bulges or internal loops within the helix. For these motifs, identifying their locations amounts to simply searching the dot-bracket data for the exact dot-bracket sequence defined for each hairpin size. For example, a regular hairpin with stem length 4 and loop length 4 has dot-bracket sequence (“((((....))))”); a regular hairpin with stem length 7 and loop length 5 has dot-bracket sequence (“(((((((.....)))))))”). As before, loops which are involved in pseudo-knotted base pairing are treated as unpaired loops for the purpose of hairpin identification.

#### Regular hairpins with or without bulges

Identifying locations of regular hairpins that may also have one or two bulges was performed similarly to the identification procedure used for regular hairpins. However, due to the increased flexibility of dot-bracket sequences and combinatorial explosion of qualified motifs when allowing for bulges, we used a regular expression scheme to perform the search. The regular expression has the form “({2,10}.{0,5}({3,10}.{3,MAXLOOP}){3,10}.{0,5}){2,10}” where MAXLOOP is the maximum loop length to include in the search. As the flexibility to allow for bulges in our regular expression necessitates the inclusion of stems possibly longer than 15 nt, any constructed structure patterns with a stem longer than 15 nt through were manually discarded prior to the search. As before, loops which are involved in pseudo-knotted base pairing are treated as unpaired loops for the purpose of hairpin identification.

### Representative RNA Structures from STRAND

To more comprehensively assess the properties of hairpins within RNA structures, we compiled a dataset of quality reference structures from the RNA Secondary Structure and Statistical Analysis Database (RNA STRAND) (59). This database houses 4666 high-quality RNA structures as determined from NMR, X-ray crystallography, or comparative sequence analysis. For our work, we pruned the number of structures significantly (to 797 structures) to account for unequal representation of RNA types within the database (for example, overrepresentation of ribosomal RNA structures). This pruning was achieved by sampling a defined number of structures from each RNA type in the database. The total numbers of original structures within each RNA type, as well as the corresponding numbers of RNA structures sampled, are given in Supplementary Table S1. A simple visualization of the fraction of (1) transcripts, (2) nucleotides, and (3) hairpins in the data coming from each RNA class is given in Supplementary Figure S1.

### Discretized Observation Model (DOM)

The discretized observation model serves as an alternative approach for describing the probabilities of a particular state (unpaired/paired) to yield a particular reactivity value (state emission distributions). Typically, the emission distributions are modeled as continuous distributions, as is the case when *patteRNA* uses a GMM of reactivity. However, the DOM framework instead discretizes reactivities based on percentiles, then constructs probability mass functions (PMFs) over the discrete reactivity classes for the two pairing states. The state PMFs are then learned in an unsupervised fashion by coupling the emission model to an HMM and performing expectation-maximization (EM) optimization of parameters, analogously to the original GMM implementation. Also analogous to the GMM’s number of Gaussian kernels, the resolution of bins used in the DOM is gradually increased until an optimal model is reached via a minimum in Bayesian information criteria (BIC) (58). Typically, 7–10 bins are deemed optimal.

A more complete description of the mathematical formulation behind the DOM, including initialization and M-step parameter updating, is available in **Supplementary Material**.

### Scoring with *patteRNA*

*patteRNA* mines structural elements as represented in dot-bracket notation. In the context of *patteRNA*, this representation of a structure is referred to as a target motif. To mine for a motif, *patteRNA* first encodes the structure as a sequence of pairing states (0: unpaired, 1: paired), called the target path. Then, all possible locations in the data are scored for the presence of the target path. With sequence constraints enforced, this amounts to all sites in an RNA where the nucleotide sequence permits folding of the target motif via Watson-Crick and Wobble base pairs. Sequence constraints can also be disabled, and in such situations all windows of length equal to the length of target motif are considered. (i.e., a full sliding window approach). Regardless of sequence constraints, the *patteRNA* score for a site (a window of length *n* beginning at nucleotide *m*) is computed as the log ratio of joint probabilities between the target path and its inverse path (i.e., the opposite binary sequence) (57). More specifically,

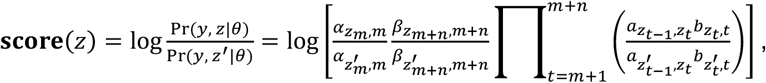

where *y* is the reactivity profile at a site, *z* is the target path, *z*′ is the inverse path, *θ* represents the parameters of a trained GMM/DOM-HMM model, *a*_*i,j*_ is the transition probability from state *i* to state *j*, *b*_*i,t*_ is the emission likelihood for state *i* at nucleotide *t*, and *α_i,t_* and *α_i,t_* are the forward and backward probabilities for state *i* at nucleotide *t*, respectively. A score of zero indicates the target path and inverse path are equally likely, and a positive score indicates the target path is more likely (and vice versa). Locations with the highest scores are subsequently deemed most likely to harbor the target motif.

To facilitate the comparative analysis of scores between different motifs and datasets, scores were further processed into *c*-scores by normalizing against a null distribution of scores estimated via sampling of scores from locations which violate the sequence compatibility necessary for the motif’s base pairs (and therefore can be presumed to not harbor the target motif) (58). Example SP data with real *patteRNA* scores superimposed is illustrated in **Figure 1A**. To mine data for the collection of all regular hairpins, the flag “--hairpins” was used when calling *patteRNA*, which automatically constructs the search space describing such motifs.

### Computation of Statistical Performance Metrics

The accuracy of *patteRNA* to detect motifs is primarily assessed through the receiver operating characteristic (ROC) and precision-recall (PR) curves. These curves were computed by varying a theoretical *c*-score threshold between called positives and negatives and, at each threshold, computing the true-positive rate (TPR/recall), false positive rate (FPR), and precision (also referred to as positive predictive value, PPV). A site is deemed a positive if all base pairs in the target motif are also present in the corresponding location of the reference structure. These performance profiles are then visualized (ROC: FPR vs. TPR, PR: TPR vs. PPV) and summarized using the area under the curve (AUC) of the ROC and average precision (AP) of the precision-recall curve. The Scikit-learn Python module (v0.24) was utilized to perform these computations.

### Data

Details about the datasets used throughout this study are compiled in Table 1.

**Table 1.** Summary of datasets used throughout this study.

| Dataset Name | Description | Size | References |
| --- | --- | --- | --- |
| Weeks set | 22 well-studied RNAs with reference structures and high-quality SHAPE data | 11,070 nt | (18, 20, 41, 57, 58) |
| STRAND data | 797 diverse RNAs with experimentally determined structures (via NMR, crystallography, or comparative sequence analysis) [no probing data] | 276,290 nt | This work, (59) |
| Manfredonia data | SARS-CoV-2 genome probed by:<br>• *In vitro* DMS-MaP<br>• *In vitro* SHAPE-MaP<br>• *In vivo* SHAPE-MaP | 3 x 29,903 nt | (33) |
| Mustoe data | 194 *E. coli* mRNA transcripts probed by SHAPE across three conditions (each condition is the average of two replicates)<br>• Cellfree (*in vitro*)<br>• Incell (*in vivo*)<br>• Kasugamycin (*in vivo* + 10 mg/mL kasugamycin) | 3 x 442,421 nt | (30) |
| Corley data | *In vivo* and *in vitro* icSHAPE data (as well as fSHAPE data, not included in the dataset size) for RNA transcripts in two human cell lines: K562 and HepG2 (each condition is the average of two replicates) | 2 x 40.8 million nt (K562)<br>2 x 35.4 million nt (HepG2) | (32) |

### Simulated Datasets and Benchmarks

We generated simulated data for RNAs in the Weeks set by sampling reactivities according to various state distributions schemes (see **Table 2**). 50 replicates of each scheme were generated for the performance benchmarks using in-house Python scripts. *patteRNA* was then used the train and mine the replicates for regular hairpins using the “patteRNA ${SHAPE} ${OUTPUT} -f ${FASTA} [--GMM or --DOM] --hairpins” command. The “-l” flag was added to use log-transformed data where applicable; training was performed independently for each replicate. Overall performance for a scheme was summarized as the mean of average precisions for the 50 replicates.

**Table 2.** Parameters of state distributions used to generate artificial data on the Weeks set. GEV: generalized extreme value.

| Scheme Name | Paired Distribution | Unpaired Distribution |
| --- | --- | --- |
| Heitsch distributions (63) | *Helix-end:*<br>GEV($\mu = 0.09$, $\sigma = 0.114$, $\xi = -0.821$)<br>*Stacked:*<br>GEV($\mu = 0.04$, $\sigma = 0.040$, $\xi = -0.763$) | Exponential distribution with $\lambda = 1.468$ |
| Gaussian / Gaussian (poor) | Gaussian distribution with $\mu = 0$, $\sigma = 1$ | Gaussian distribution with $\mu = 0.5$, $\sigma = 1$ |
| Gaussian / Gaussian (medium) | Gaussian distribution with $\mu = 0$, $\sigma = 1$ | Gaussian distribution with $\mu = 1$, $\sigma = 1$ |
| Gaussian / Gaussian (high) | Gaussian distribution with $\mu = 0$, $\sigma = 1$ | Gaussian distribution with $\mu = 2$, $\sigma = 1$ |
| Exponential / Gaussian | Exponential distribution with $\lambda = 2$ | Gaussian distribution with $\mu = 2$, $\sigma = 1$ |
| Exponential / Exponential | Exponential distribution with $\lambda = 2$ | Exponential distribution with $\lambda = 1/2$ |

### Hairpin Mining Performance of NNTM Partition Function Approach

We benchmarked the performance of partition function approaches to detect hairpins in the Weeks set by using the “RNAsubopt” command from ViennaRNA to generate 1000 structures for each transcript in the Weeks set, using that transcript’s SHAPE data as soft constraints (“RNAsubopt -p 1000 --shape ${SHAPE_FILE} < ${SEQUENCE}”). For each hairpin in the generated structural ensemble, a “score” was assigned as the fraction of structures in the structural ensemble which contain the base pairs comprising that hairpin. Predicted hairpins and their scores were organized into a single list which was then processed into a receiver operating characteristic and precision-recall curve as done for *patteRNA*’s predicted hairpins (see *Computation of Statistical Performance Metrics*).

### Posterior Pairing Probabilities

*patteRNA* computes pairing probabilities as described (57). Briefly, a parameterized GMM-HMM or DOM-HMM model is utilized to compute emission likelihoods for each nucleotide, followed by the forward and backward probabilities via the forward-backward algorithm. Posteriors are then computed as the product of the forward and backward probabilities and appropriately scaled such that *P*(paired) + *P*(unpaired) = 1 for each nucleotide.

### Hairpin-Driven Structure Level (HDSL)

The hairpin-driven structure level (HDSL) is a nucleotide-wise measure quantifying the local level of structure from SP data. HDSL is initialized using posterior probabilities to be paired as computed by *patteRNA*. Then, the profile is augmented using high-scoring hairpins from *patteRNA*. For each detected hairpin with *c*-score greater than 0.5, the value 0.2 · (*c*-score – 0.5) is added to the profile at all nucleotides covered by the hairpin. After profile augmentation, profile smoothing is achieved via a 5 nt sliding-window mean followed by a 15 nt sliding-window median to give the final HDSL profile. Analogous approaches using just a sliding mean or just a sliding median were also tested, but we found that the best results were obtained when coupling the two summary statistics together.

A flow chart illustrating the flow of information as handled by *patteRNA*, including the relationship between HDSL and the training and scoring phases, is included as **Figure 2**.

**Figure 2.**
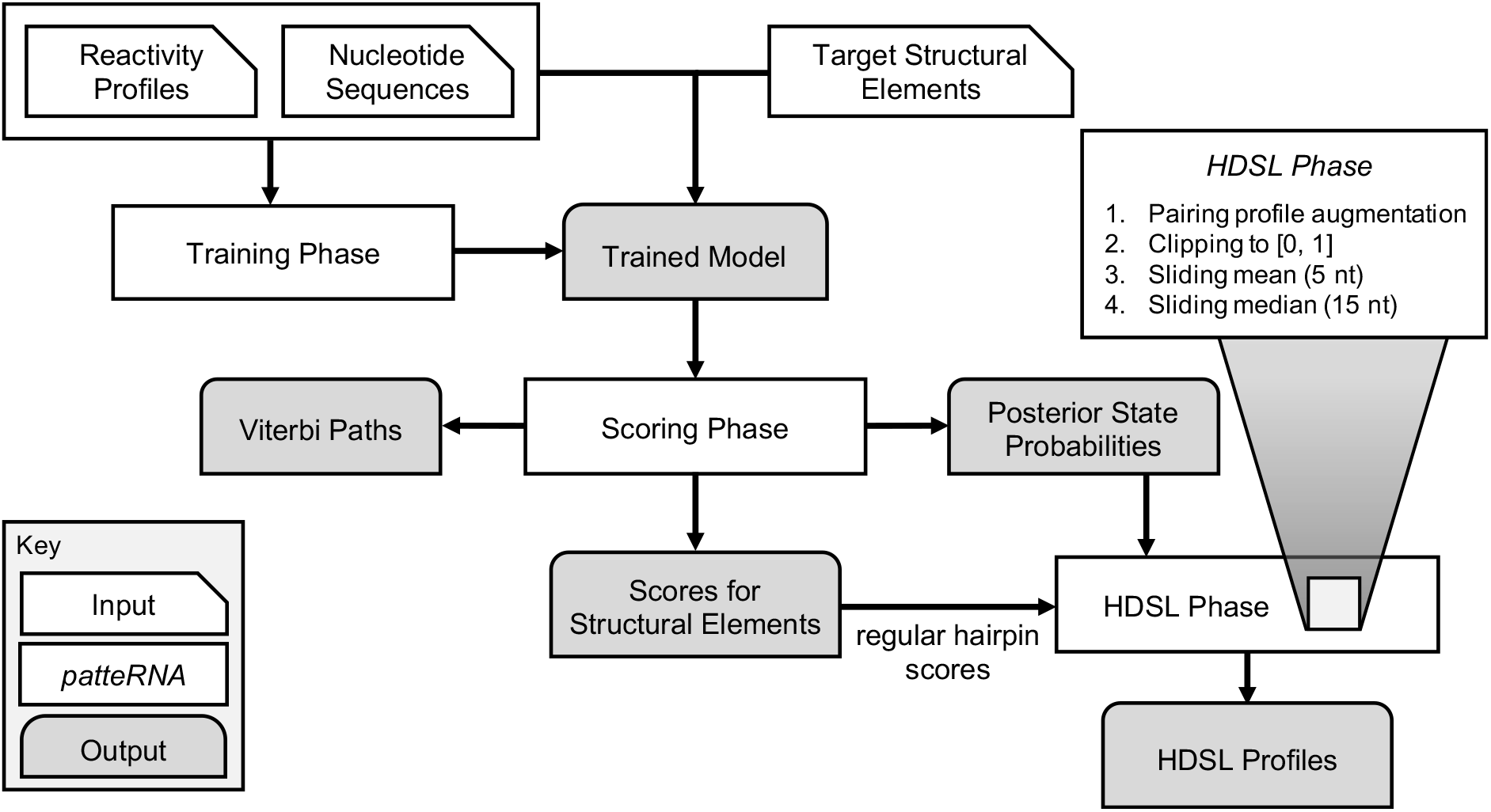
Overall flow of data and computing behind *patteRNA* and hairpin-derived structure level (HDSL). The measure is initialized as the pairing probability profiles, which are then augmented by boosting values at sites covered by highly scored hairpins (see Methods). The subsequent profile is clipped to the interval [0, 1] and local smoothing is achieved with sliding window mean and sliding window median approaches with windows of size of 5 nt and 15 nt, respectively.

### Averaging and Integrating HDSL over mRNA Coding Sequences

We delineated the regions surrounding the 432 genes in the Mustoe data into 4 groups: (1) start site; ±30 nt around <u>A</u>UG, (2) 5’UTR; −70 to −31 nt from <u>A</u>UG, (3) 3’ UTR; +1 to +40 from STOP codon, and (4) coding sequences; +31 nt from <u>A</u>UG to the STOP codon. For the start site, 5’UTR, and 3’UTR, averages were taken at each aligned position as these groups each have a constant length. For situations where all regions might not exist for a gene, aligned HDSL profiles were included in the analysis as far as the nucleotide sequence allowed, and remaining positions were treated as missing values and omitted from subsequent averaging. For instance, if the 5’UTR was 50 nt (i.e., less than 70 nt), those 50 nt were aligned with the corresponding locations and the missing 20 nt upstream were treated as missing values. For coding sequences (which inherently have a non-constant distribution of lengths), the profiles were interpolated to a vector of length 300 to allow for aligned averaging relative to the beginning and end of the window. 99% confidence intervals were computed using the Wald formulation (mean HDSL ± 2.576 · SE).

### Local Folding Calculations

Windowed partition function calculations were performed using the “RNAfold -p” command from ViennaRNA (19). Three schemes were utilized: windows of length 3000 nt, spaced 300 nt apart; windows of length 2000 nt, spaced 150 nt apart; and windows of length 150, spaced 15 nt apart. In each case, sequences within each window were parsed using custom Python scripts and then processed sequentially with RNAfold. Only the time required to run RNAfold commands was measured in timing benchmarks (no integration of windowed outputs or post-processing were accounted for). RNALfold benchmarks were performed using the default arguments of the command to process all sequences in the Corley data sequentially. All timing comparisons in this study were performed on an AMD Ryzen 9 5900X CPU running Ubuntu 20.04 LTS.

### *patteRNA* Training and Scoring

Unless otherwise noted, all *patteRNA* analyses were performed with default training parameters (KL divergence for training set: *D*_KL_ = 0.01, convergence criterion *ε* = 0.0001, automatic determination of model complexity, *k*, via Bayesian information criteria) (58). With the exception of benchmarks investigating the effect of log-transforming data, log-transformed data were always used when using a GMM and non-transformed data were used when using a DOM. Scoring for regular hairpins was achieved using the “--hairpins” flag and computation of HDSL profiles was achieved with the “--hdsl” flag.

## RESULTS

### Overview of *patteRNA* Mining

To mine structure elements from SP data, *patteRNA* first learns the statistical properties of the data via the training phase. The purpose of this procedure is to estimate the distributions of reactivities associated with paired and unpaired nucleotides, respectively. Training is unsupervised and has been shown to accommodate diverse data distributions (see Ledda et al. (57) for a complete description). With the dataset characterized via its statistical model, *patteRNA* can then mine for structural motifs.

**Figure 1A** demonstrates key concepts related to *patteRNA*’s motif mining. When mining a particular structural element (i.e., the target), sites which satisfy the sequence constraints necessary for the target’s secondary structure are scored for their probing data’s consistency with its pairing state sequence (57, 58). Sites which do not satisfy sequence constraints can also be scored, however these sites are almost certainly all negatives and can therefore be discarded (the only exception being the possibility of non-canonical base pairs). Sites which harbor the target motif presumably have SP data consistent with the desired state sequence and therefore score highly. *patteRNA*’s overall objective is to identify sites harboring particular structural elements, such as hairpins, as accurately as possible.

### Hairpins Comprise a Significant Portion of Structural Elements

To assess the plausibility of a hairpin-centric approach in making general assessments of structure, we examined a diverse dataset of 22 RNAs with known structures (~10,000 nt) (57) to quantify the distribution of hairpins present as well as the proportion of base pairs contained within hairpins. We refer to this dataset as "the Weeks set." Analyzing the 278 distinct hairpins in the Weeks set reveals that a majority fall within a narrow range of stem and loop lengths (**Figure 1B**). Specifically, hairpins most frequently have loop lengths between 3 and 10 nt, and stem lengths 15 nt or less. In other words, although their properties are diverse, there is a range of stem and loop sizes which represents a majority of hairpins (83%). Later in the study will we leverage these characteristic properties to focus our searches on this most representative subset of hairpins.

Our results also illustrate that hairpins comprise a large fraction of structural elements. We first focused on hairpins with no bulges or internal loops (i.e., unpaired stretches flanked by some number of base pairs), which we call regular hairpins, and found that around 35% of paired nucleotides reside in such structures (**Figure 1C**). If you also consider hairpins with up to two bulges each with length up to 5 nt, this coverage increases to over 50%. This suggests that, although hairpins are only a subset of RNA structural elements, they are indeed the most prevalent, and therefore identifying them in SP data could provide a strong quantification of general structural trends.

Understanding that the Weeks set is a small sample of structures to draw conclusions from, we repeated this hairpin counting and quantification on a diverse set of 797 reference structures from the STRAND database (59) representing a more complete profile of RNA structure properties. The distribution of hairpins in this dataset is shown in **Supplementary Figure S2** and recapitulates the observations from the Weeks set. In fact, the STRAND data suggests that regular hairpins specifically comprise a slightly larger fraction (40%) of structural elements than is seen in the Weeks set (35%).

One can further expand the definition of a hairpin to also include the associated stems that extend from a hairpin element up to the first nucleotide that base-pairs outside of the nested context of this element (see **Supplementary Figure S3** for examples). We refer to these helices as external stems and note that such motifs are prevalent in structured RNAs. **Figure 1C** shows that relaxing the definition of a hairpin to include external stems leads to over 80% coverage of paired nucleotides. Although such elements are beyond the scope of the analysis that follows, this high coverage indicates that a large majority of RNA structure can be represented as motifs with local base pairing. Moreover, it’s important to note that virtually all types of canonical RNA structure motifs necessarily exist in the context of hairpin elements—internal stems, multibranch junctions, etc., only exist in the presence of hierarchical domains which all terminate in a hairpin-like fashion.

### Simplified Reactivity Model Improves Accuracy of Motif Detection

In an attempt to improve *patteRNA*’s performance, we investigated alternative statistical models of reactivity and their downstream effects on scoring accuracy. While the GMM approach performs well, especially at the task of approximating the underlying state distributions, we encountered issues in motif scoring. Namely, reactivities from the tails of the overall data distribution would be strongly predicted to be paired or unpaired. This isn’t an inherent problem, as the most extreme reactivities should theoretically be the best candidates for confident prediction. However, these reactivities present problems during scoring as they have the propensity to dominate the score for sites they fall into. In other words, a single extreme reactivity consistent with the target state sequence could yield a high score for a site, even if data within that site is otherwise inconsistent with the target (and vice versa). Generally speaking, for SP data such as SHAPE, the most extreme reactivities are only about 3-5 times more likely to be in one state over the other (60), yet the GMM often arrives at likelihood ratios 10 or 100 times larger than this empirical ratio. Such predictions have negative consequences on the interpretation of scores.

Motivated by these issues, we devised a simplified framework for unsupervised learning of the state reactivity distributions. It entails a discretized observation model (DOM) which substitutes for the GMM component of the statistical model (i.e., the emission probabilities), resulting in a DOM-HMM model of SP data. The DOM entails modeling reactivities as a discrete distribution where they are binned into classes based on percentiles. During training, pseudo-counts are estimated for each class (E-step) and then utilized in the M-step to infer the discrete reactivity distribution for paired and unpaired states. A schematical comparison of the GMM and DOM approaches is shown in **Figure 3A** (see Methods and **Supplementary Material** for a complete mathematical formulation).

**Figure 3.**
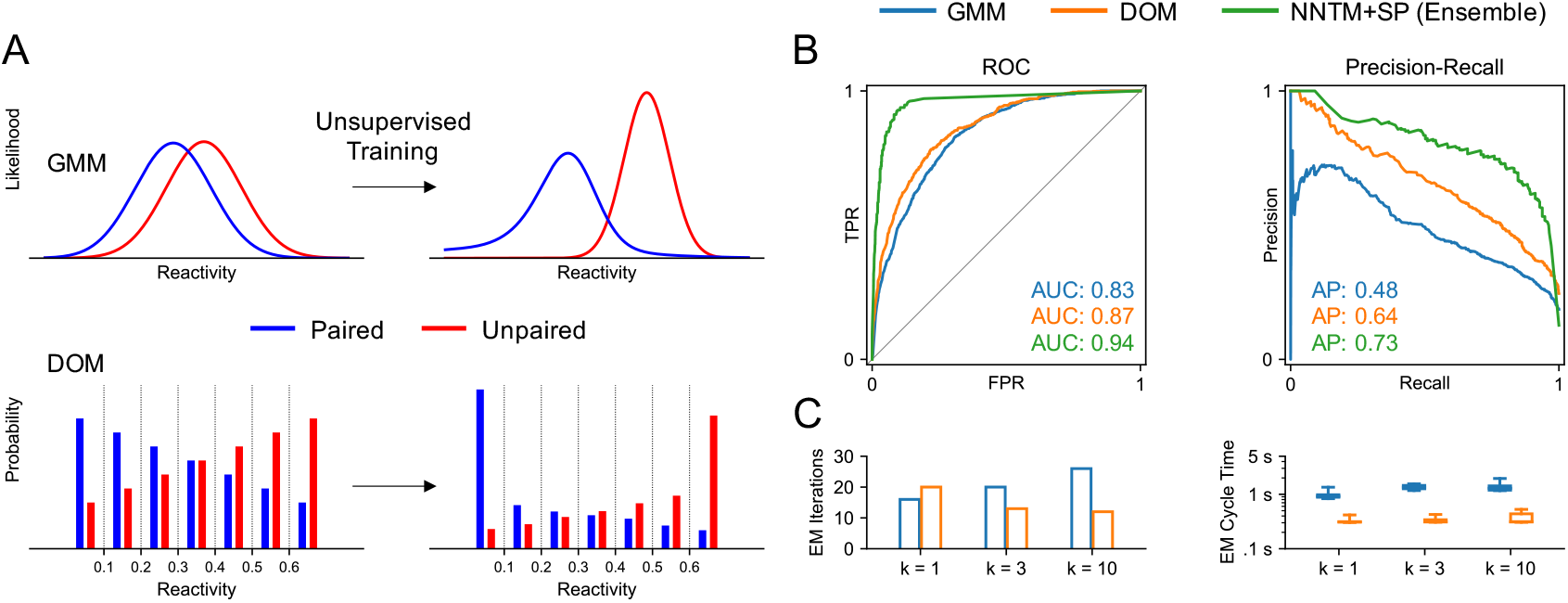
A discretized observation model (DOM) of reactivity improves hairpin detection precision when compared to a Gaussian mixture model (GMM). (A) Schematic illustration of GMM and DOM approaches in the content of *patteRNA*’s unsupervised learning scheme. The DOM is founded upon a percentile-based discretization of reactivities which yields a discrete emission probability scheme. The discretization scheme it itself optimized during training based on Bayesian information criteria (BIC) of models using progressively smaller bins. (B) Receiver operating characteristic curves and precision-recall curves when mining regular hairpins in a reference dataset (“the Weeks set,” see text) with *patteRNA* using either GMM (blue) or DOM (orange) approaches, or when using data-driven NNTM-based folding (green). (C) Timing benchmarks of unsupervised training via GMM and DOM on the Weeks set. Shown are the number of EM iterations required for convergence on the Weeks set and time required for a single EM iteration. 5 repetitions were used when measuring EM cycle times.

We benchmarked the capacity of *patteRNA* to identify regular hairpins in the Weeks set via the GMM and DOM. We assessed their discriminatory power primarily via the receiver operating characteristic (ROC) and precision-recall curve (PRC), which are shown in **Figure 3B**. Our results indicate that the DOM approach improves both the area-under-the-curve (AUC) of the ROC and the average precision (AP) of the PRC. Although the improvement to AUC appears minor, average precision was increased from 0.48 with a GMM to 0.64 with a DOM. Precision is a crucial performance metric in structure motif mining where the vast majority of scored sites are negatives (even with sequence constraints applied), so the improvements seen in the DOM are important through this perspective. Notably, precision at the highest scores is much better in the DOM compared to the GMM, which is susceptible to numerous negatives at the highest hairpin scores despite decent precision at moderate scores. This is evidenced by the large fluctuations in precision at low levels of recall for the GMM (see the top left of precision-recall plot in **Figure 3B**). The DOM approach, on the other hand, is far more reliable for returning positive hits at the highest scores. **Figure 3B** also includes a benchmark for data-directed NNTM folding algorithms which shows that *patteRNA* is, although improved via the DOM, generally unable to match the precision of RNA folding. Notably, NNTM folding was performed with an ensemble-based approach, which, although much slower, outperforms a single MFE calculation (57).

Importantly, the presented results show overall performance on the collection of all regular hairpins, which is comprised predominantly by motifs with shorter stems. Shorter stems present a challenge to *patteRNA*, as fewer base pairs render sequence constraints less effective in controlling the number of negative sites considered in the analysis. When comparing performance on individual motifs, however, we find that *patteRNA* matches the precision of NNTM-ensemble methods for longer stems. In some cases, such as hairpins with stem length 6 and loop length 7, it even surpasses the performance of the NNTM approach (see **Supplementary Figure S4**). We also observe a universal trend for the DOM to outperform the GMM at the motif-level, further validating its superior performance.

Not only does the DOM improve precision, but the model itself is described by fewer parameters and trains faster than a GMM. As seen in **Figure 3C**, faster training is achieved in two distinct ways. First, the DOM generally requires fewer EM iterations to converge. Second, EM iterations are significantly faster. The latter is presumably due to the DOM’s simpler M-step formulation, which reduces to simple counting as opposed to the GMM which requires multiplication and squaring to update the means and variances of each Gaussian kernel.

Given the rapidly evolving field of structure probing and disparate statistical properties of SP datasets (47), we also investigated whether the benefits from the DOM generalize to other data distributions. Different probes have different quality (47, 61), different conditions yield different quality (47), and the quality of probes is constantly improving (62); therefore, adaptability of methods is crucial. Benchmark datasets like the Weeks set are not currently available for the plethora of probes used, so we resorted to simulations. We constructed several artificial datasets and benchmarked *patteRNA*’s performance via the GMM or DOM approaches. We sampled reactivities for the underlying structures in the Weeks set according to various state distributions, including empirically-fitted distribution models from Sükösd et al. (63), referred to as the Heitsch distributions, as well as a collection of mock distributions with varying classification power (i.e., various degrees of separation between the state distributions). For each scheme, 50 replicates were created, and we benchmarked performance against both the regular and log-transformed data. We note that the fidelity of the GMM is dependent on the Gaussianity of the data, presenting a weakness of this approach as the decision to log-transform can have a major impact on scoring efficacy.

The results of the benchmarks are shown in **Table 3**. Generally speaking, the DOM matches or exceeds the performance of the GMM. Depending on the data properties, the DOM’s performance gain ranges from minute to transformative. In only one of the benchmarks did the GMM outperform the DOM (poor quality Gaussian / Gaussian data), and only by a small margin. This specific outcome might be explained by the DOM’s simplification of SP data which effectively clips extreme reactivities when discretizing the data. In datasets of poor quality, the most extreme reactivities likely provide the only opportunity for reliable inference on pairing state, so it’s possible that the relatively coarse discretization scheme reduces the information content of the data. Regardless, it’s worth noting that data of such poor quality is uncommon, especially in light of on-going improvements to experimental protocols and probe quality (8, 10, 62, 64). Our results also demonstrate the adaptability of the DOM and its robustness to non-Gaussian data, which render the method broadly applicable. When using the DOM, log-transforming is largely irrelevant to model performance, as the discretization scheme is founded on data percentiles. The lone exception to this rule is when handling reactivities below zero, which are necessarily binned together if data is log-transformed.

**Table 3.**
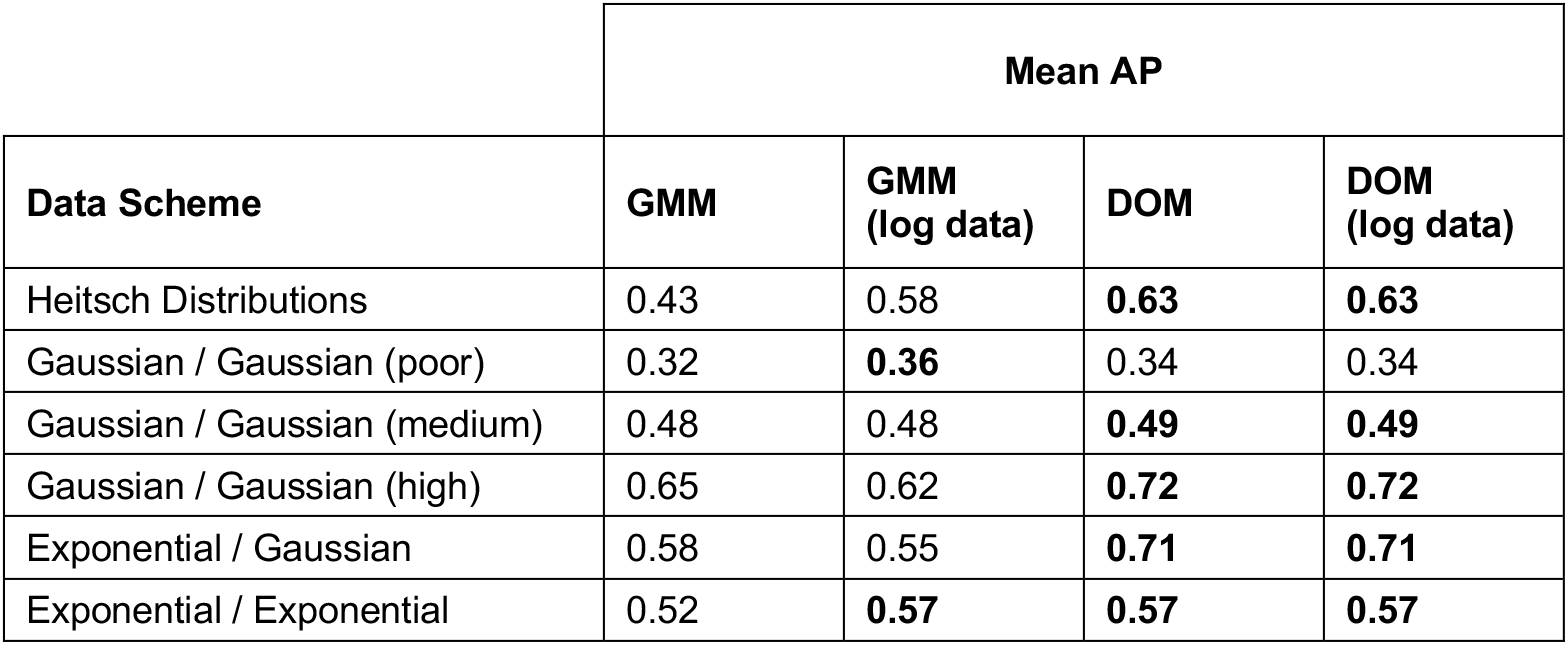
Average precisions of *patteRNA* for hairpin mining when utilizing a Gaussian mixture model (GMM) or discretized observation model (DOM) of reactivity against various artificial data schemes. For all benchmarks, average precision was averaged over 10 replicates. Bold entries highlight the best performing approaches for each scheme. AP: average precision.

Overall, these results demonstrate the benefit of the DOM approach in more efficiently and effectively mining structures from SP data. Note, however, that the GMM still provides a specific utility when one’s objective is to arrive at continuous models of the state reactivity distributions (e.g., to use for simulations, or for data inspection). *patteRNA* includes both implementations such that the respective approach can be used depending on the intended use-case.

### Summarizing Structuredness in RNAs from Hairpin Detection

As hairpins comprise a large fraction of structural elements, we sought to utilize *patteRNA* to quantitatively summarize local “structuredness.” Due to the plethora of cellular processes affected by RNA structures, there are numerous contexts in which summarizing local structure is important. To name a few examples, one might wish to find structural domains and druggable pockets in viral genomes (29, 33, 65), quantify connections between mRNA structure and gene regulation (43, 48, 57, 66–68), identify transcriptome-wide where RNA is differentially affected by particular stimuli (32, 69), or compare structure between conditions and/or logical regions of genomes (1, 30, 70). The most popular approach for quantifying structuredness relies on a combination of two metrics: local reactivity and local Shannon entropy. Local reactivity is generally computed via a rolling mean or median with windows ranging 25-500 nt, while local Shannon entropy derives from base pairing probabilities computed via NNTM folding routines. The combination of these two metrics yields regions which are largely unreactive (i.e., base paired) and stable (i.e., tending to adopt one conformation). We note that each metric by itself is generally insufficient in this context, as low reactivity regions sometimes include regions which see multiple competing conformations (but are nevertheless highly paired), and low Shannon entropy can also be observed for regions which are preferentially single stranded.

To integrate *patteRNA*’s results into a quantification of structuredness, we propose a nucleotide-wise measure we term the hairpin-derived structure level, or HDSL. At the highest level, HDSL combines *patteRNA*’s computed base pairing probabilities with information from hairpin searches. This allows us to consider the locations of stable hairpins in addition to the overall pairing propensity of regions, the former of which typically does not account for all structured regions (e.g., external stems, stems with numerous bulges, or stems with non-canonical base-pairing). Briefly, the posterior pairing probabilities are used as a starting point. They are then amplified at nucleotides covered by highly scored hairpins, depending on the hairpin score—the higher a hairpin is scored, the larger the boost. Next, the profile is clipped to [0, 1] and locally smoothed by taking a 5 nt rolling mean followed by a 15 nt rolling median (see Methods for a complete description). We explored the properties of HDSL and validated its utility as an indicator of local structure by applying it to three recent datasets that were previously used to assess local structuredness in diverse contexts.

### Trends in Detected Hairpins Recapitulate Known mRNA Dynamics in *E. coli*

We analyzed the set of 197 mRNA transcripts (comprising 432 genes) in *E. coli* probed *in vitro*, *in vivo*, and *in vivo* + kasugamycin with SHAPE-MaP by Mustoe et al. (30). In addition to Mustoe et al.’s analysis, previous studies have demonstrated that mRNAs fold differentially in cells compared to *in vitro* (50, 66, 70, 71). *In vivo* mRNAs have been observed to be less structured than their *in vitro* counterparts, with the magnitude of structural changes correlated with translation (31, 72). These effects have been observed most strongly in the context of the 5’UTR and CDS of highly expressed genes. Conversely, structural changes have also been observed around the 3’UTR, but evidence demonstrating both a decrease (71) and increase (72) in structures has been published in the literature, possibly correlating to the degree of post-transcriptional regulation of transcript decay (72). We applied HDSL to Mustoe et al.’s data and investigated to what degree our measure reveals structural changes along mRNA transcripts in a prokaryotic organism like *E. coli*.

The results of our analysis are compiled in **Figure 4**. In **Figure 4A**, we compare averaged HDSL profiles over the 432 genes included in the study between *in vitro* and *in vivo* conditions. The averaged HDSL profiles are delineated into 3 groups: nucleotides near the start site (AUG ± 30 nt), nucleotides within the coding sequence (at least 31 nt downstream of AUG), and nucleotides in UTRs (5’UTR: 31–70 nt upstream of AUG; 3’UTR: first 40 nt after STOP). Our results demonstrate that, as expected, UTRs are generally the most structured regions of the transcripts. They also show a strong intrinsic effect for mRNA to be relatively less structured around the start codon in both conditions. Moreover, *in vivo* data show that factors in this condition work to further unfold structures around the start site, as HDSL is significantly lower around the start codon *in vivo* than *in vitro*. Additionally, we did not detect a strong signal for structures in coding sequences to be de-structured overall when accounting for the region around the start codon separately. In contrast, Mustoe et al.’s analysis, which did not consider start sites separately (i.e., considered only CDS versus non-CDS), concluded that coding sequences are relatively unstructured in cells based on a slight increase in reactivities *in vivo* versus *in vitro* for nucleotides in CDS. The subtle unfolding effects observed in Mustoe et al. may be partially attributed to localized effects around the start codon observed here. Notably, the specific relevance of structure around this region of mRNA transcripts has been observed and recognized as important in several other studies on organisms of varying genetic complexity (45, 46, 50, 73).

**Figure 4.**
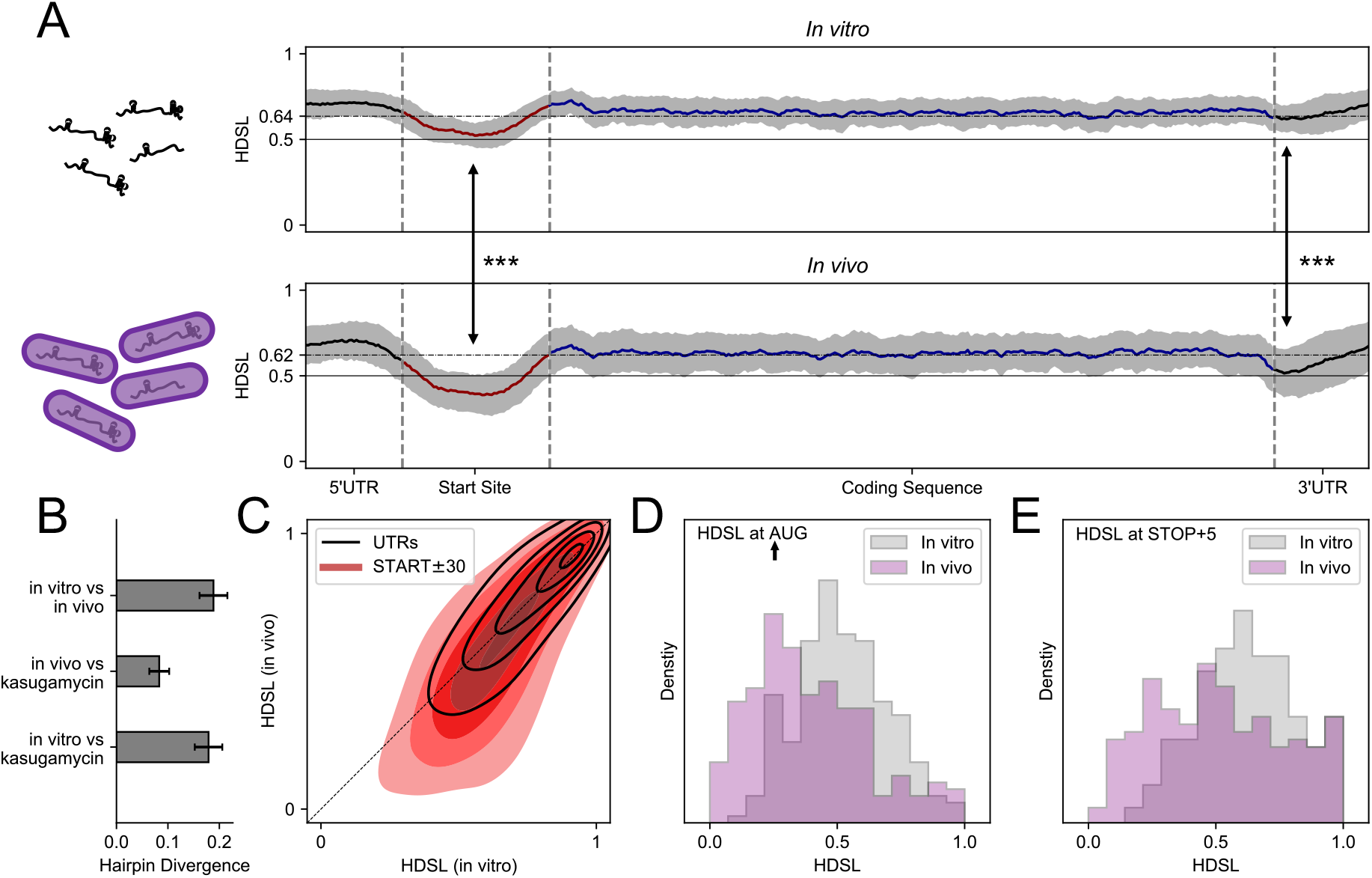
Hairpin-derived structure level (HDSL) demonstrates regional differences in structure changes between *in vivo* and *in vitro* structures for mRNA transcripts in *E. coli* (probed by Mustoe et al. (30)). (A) Averaged HDSL profiles across all genes (*N* = 432) for nucleotides around the start codon (±30 nt, red), within the coding sequence (AUG+31 to STOP), and 5’/3’UTRs (black). Grey area indicates the 99% CI of mean HDSL. Dot-dashed lines indicate mean HDSL over all nucleotides. (B) Hairpin divergence (fraction of *patteRNA*-detected hairpins unique to one condition) for the three pairwise comparisons between *in vivo*, *in vitro*, and *in vivo* + kasugamycin conditions. Error bars represent the exact binomial (Clopper-Pearson) 99% CI. (C) 2D density plot of HDSL between the two conditions shown in (A) indicates a bias for weakly structured regions *in vitro* to become more unstructured *in vivo*. (D) Histograms of HDSL at the adenosine of start codons for both conditions in (A). (E) Histograms of HDSL at the fifth nucleotide after the STOP codon for both conditions in (A). Arrows indicate Wilcoxon signed-rank tests for mean HDSL at the noted positions to be equal in both conditions. *** indicates *p* < 1 × 10^−60^.

To further substantiate the effects we observed, we checked the similarity of *patteRNA*’s detected hairpins for each pairwise comparison of the three conditions included in the original study. Ideally, in the absence of significant structural remodeling between two conditions, we expect to find the same hairpins in both. On the other hand, if two conditions are substantially different, we expect to see larger differences in the hairpins detected by *patteRNA*. Searching for the aforementioned set of regular hairpins (see *Hairpins Comprise a Significant Portion of Structural Elements*) and using a *c*-score threshold of 1 to indicate a “detected” hairpin, we computed the fraction of hairpins reproducible in both conditions of each comparison (**Figure 4B**). We see that *in vivo* and *in vivo* + kasugamycin have the highest level of hairpin conservation (less than 10% of detected hairpins are not present in both conditions, meaning >90% similarity in detected hairpins). This high similarity serves as a basic quality control measure, as the *in vivo* + kasugamycin condition, although affected by changes to translation initiation, is nevertheless highly similar to the *in vivo* condition. On the contrary, comparing *in vivo* to *in vitro* data shows that 20% of detected hairpins are unique to one condition. The very high level of similarity between *in vivo* and *in vivo* + kasugamycin reaffirms that the differences observed in **Figure 4A** between *in vivo* and *in vitro* reflect real differential effects, rather than the impact of biological variation or artifacts from *patteRNA*’s imperfect hairpin detection scheme.

To further investigate the differences between the conditions around start codons, we visualized the condition-wise correlation of HDSL for all nucleotides within this region (**Figure 4C**). We detected a tendency in this area for the most structured regions *in vitro* to remain structured *in vivo* (see top right of distribution, which is tightly concentrated around the diagonal). The density of HDSL in **Figure 4C** does reveal a tendency for HDSL to be reduced in the *in vivo* condition, but mostly for regions with moderate HDSL *in vitro*. Thus, the overall de-structuring effect from **Figure 4A** appears to be driven by unfolding of moderately structured regions. **Figure 4D** compares the HDSL distribution between *in vitro* and *in vivo* at the adenosine residue of the start codon. There is a noticeable reduction in HDSL in the *in vivo* condition (*p* < 1 × 10^−60^, Wilcoxon signed-rank test), presumably driven by translation and possibly other cellular effects destabilizing mRNA structure, as discussed above. There is also a noticeable reduction in HDSL near the start of the 3’UTR (**Figure 4E**, *p* < 1 × 10^−60^, Wilcoxon signed-rank test), although this effect disappears on average for nucleotides farther away from the end of the coding sequence (see **Figure 4A**). Overall, our results demonstrate that HDSL can rapidly measure local structure and gives results consistent with prior analyses.

### HDSL Correlates Strongly with Structured Regions of SARS-CoV-2

To further explore the properties of HDSL, we applied it to the SARS-CoV-2 genome. Recently, multiple labs have independently probed the genome with SHAPE (33, 74) and dimethyl sulfate (DMS) (33, 75). These works have resulted in a complete structure model of the genome, highlighted by the identification of structured elements across its entire length. Here, we focus on SP data generated by Manfredonia et al. (33), which contained SHAPE data both *in vitro* and *in vivo*. Other studies either had data for only one condition or relied on DMS, which only reports reactivity for A and C nucleotides.

We first characterized the consistency of *patteRNA*’s detected hairpins with the complete structure model proposed by Manfredonia et al. We took the published structure model as ground-truth, searched for all predicted regular hairpins, and quantified the accuracy of *patteRNA* via the ROC (**Figure 5A**) and PRC (**Figure 5B**). Our results reveal good consistency between detected hairpins and hairpins in the predicted genome structure, as evidenced by AUCs around 0.88 and APs above from analyses for both conditions.

**Figure 5.**
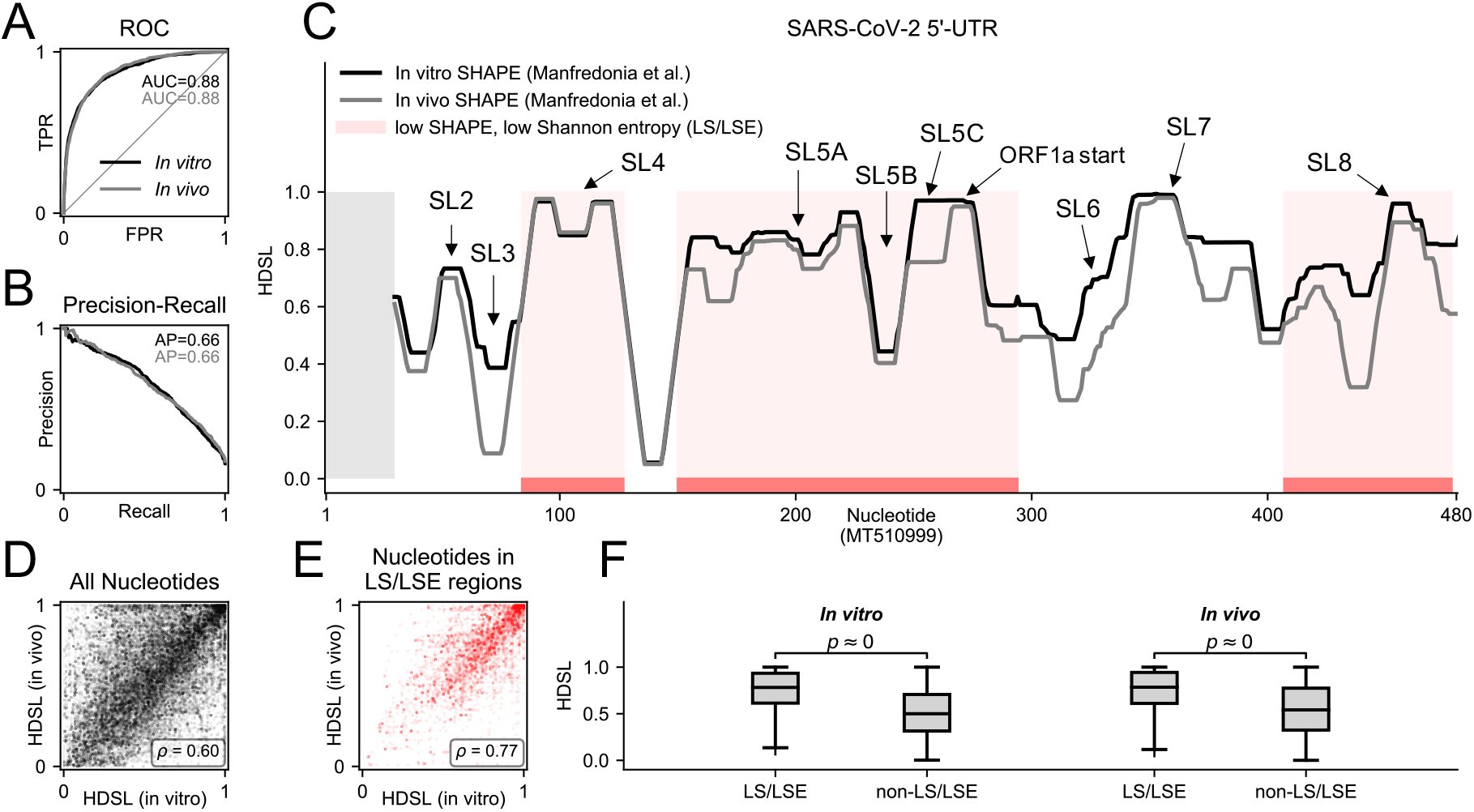
HDSL demonstrates correlated and differential structuredness between *in vitro* and *in vivo* SHAPE experiments on SARS-CoV-2 by Manfredonia et al. (33). (A, B) Receiver operating characteristic curves and precision-recall curves for *patteRNA*’s detected hairpins. (C) HDSL profiles for the 5’UTR of SARS-CoV-2 *in vitro* and *in vivo* with low SHAPE, low Shannon entropy (LS/LSE) regions (called by Manfredonia et al.) indicated in red. Grey regions indicate no data. (D) Scatterplot of HDSL *in vitro* and HDSL *in vivo* for all nucleotides of the genome. (E) Scatterplot of HDSL *in vitro* and HDSL *in vivo* for nucleotides in LS/LSE regions. (F) Boxplot comparison of HDSL profiles within LS/LSE regions and outside of them for both conditions.

Next, we used *patteRNA* to generate *in vivo* and *in vitro* HDSL profiles. Inspecting them in the 5’UTR reveals trends consistent with the currently accepted structure models (see **Figure 5C**) (33, 74–78). Namely, HDSL is high at known stable stem-loops, such as SL2, SL4, SL5A/C, SL7, and SL8. A weaker signal is found at SL6, which also shows differential structuredness between *in vitro* and *in vivo* data. Comparative analysis (79), *in vivo* RNA-RNA interactions (76), and multiple probing datasets (33, 74, 75) support the presence of this element. However, mutagenesis studies on a related coronavirus, murine coronavirus (MHV), demonstrated that disrupting this stem loop did not significantly affect virus viability (80). Given that SL6 is within of ORF1ab, it is possible that the element is transient in nature. That said, NMR experiments concluded SL6 stably forms and additionally measured a significantly larger internal loop than was predicted with *in silico* structure models (78). The internal loop, also identified as a major binding site for the N protein (81), appears to be responsible for high reactivities and the observed differential structuredness of SL6 between *in vitro* and *in vivo* data. Similarly, for SL3, although comparative sequence analysis and NNTM-based folding with *in vitro* data suggest the presence of this stem-loop, *in vivo* data does not agree with its presence (33, 74, 75). NMR investigations concluded that the stability of the element is strongly influenced by ionic conditions (78), and studies on RNA-RNA interactions suggest that this stem loop is unfolded *in vivo* to facilitate genome cyclization, as the region is involved in a long-range interaction with the 3’UTR (76). As such, differential structuredness between *in vitro* and *in vivo* conditions is consistent with current understandings of the stem-loop element. Finally, we observe relatively low HDSL for SL5B, an element confirmed via RNA-RNA interactions (76) and NMR (78). NMR studies, however, suggest that the upper part of the stem is destabilized at physiological temperatures by the presence of SL5C. The presence of a bulge and high reactivities near the apical loop of SL5B subsequently result in attenuated HDSL observations around this element, as the structure scores poorly for the regular hairpin motifs considered by *patteRNA* when summarizing structuredness. Although a complete analysis of the SARS-CoV-2 genome is beyond the scope of this study, full HDSL profiles for the two conditions are included in **Supplementary Figure S5**.

Generally speaking, there is a reasonable correlation between HDSL *in vitro* and HDSL *in vivo* (**Figure 5D**), although some deviation is expected given that *in vivo* contexts alter RNA dynamics. We also compared the properties of HDSL within Manfredonia et al.’s called “low SHAPE, low Shannon entropy” regions (regions with locally low SHAPE and Shannon entropy). Inspecting HDSL properties within these regions confirms they are characterized by very high HDSL levels, as seen in **Figure 5E** and **Figure 5F**. We investigated this association in more detail by correlating Shannon entropy with the following: SHAPE reactivity, pairing probabilities from *patteRNA*, and HDSL (**Supplementary Figure S6**). Our results show that reactivity is loosely correlated with Shannon entropy, but pairing probabilities correlate slightly better. However, HDSL shows an even stronger correlation, suggesting that it captures structuredness better than the former measures. Lastly, our results on the SARS-CoV-2 genome indicate that HDSL profiles retain sufficient resolution to capture locations of specific structural elements (e.g., individual stem-loops in the 5’UTR), boding for the plausible use of our measure to assist in more detailed analyses of regions in addition to quantifying local structuredness.

### RBPs Bind RNA at Structured Regions

Corley et al. (32) devised a novel experimental procedure called fSHAPE which can detect RNA nucleotides engaging in hydrogen bonding with RNA binding proteins (RBPs). fSHAPE works by chemically probing RNA transcripts in the presence and absence of native binding factors, then quantifying the degree of modification change between the two conditions. Nucleotides bound by RBP would presumably be more reactive in the absence of binding factors, which translates to a high fSHAPE score. Integrating fSHAPE information with standard reactivity profiles therefore allows one to examine the structural context of RBP binding sites. In this regard, Corley et al. performed icSHAPE in tandem with fSHAPE to perform such analyses transcriptome-wide on human cell lines (K562, HepG2, and HeLa). Their work showed that nucleotides with high fSHAPE scores tend to fall in areas with relatively low Shannon entropy when compared to the regions flanking them, allowing them to conclude that RBP tend to associate with RNA in the general context of structured regions. We sought to use HDSL to address the same question, namely, is there a structural context characteristic to RBP binding? To this end, we processed their icSHAPE data with *patteRNA*, mined for regular hairpins, and computed HDSL profiles. We first investigated what association exists, if any, between high fSHAPE nucleotides and pairing probabilities as computed by *patteRNA*’s DOM-HMM. Simply put, we found that nucleotides with high fSHAPE (fSHAPE > 2) are almost unanimously unpaired (**Figure 6A**), while nucleotides with lower fSHAPE follow a distribution encompassing both states yet biased towards paired states. The association of high fSHAPE with unpaired nucleotides recapitulates what Corley et al. demonstrated with pairing probabilities computed via partition function approaches.

**Figure 6.**
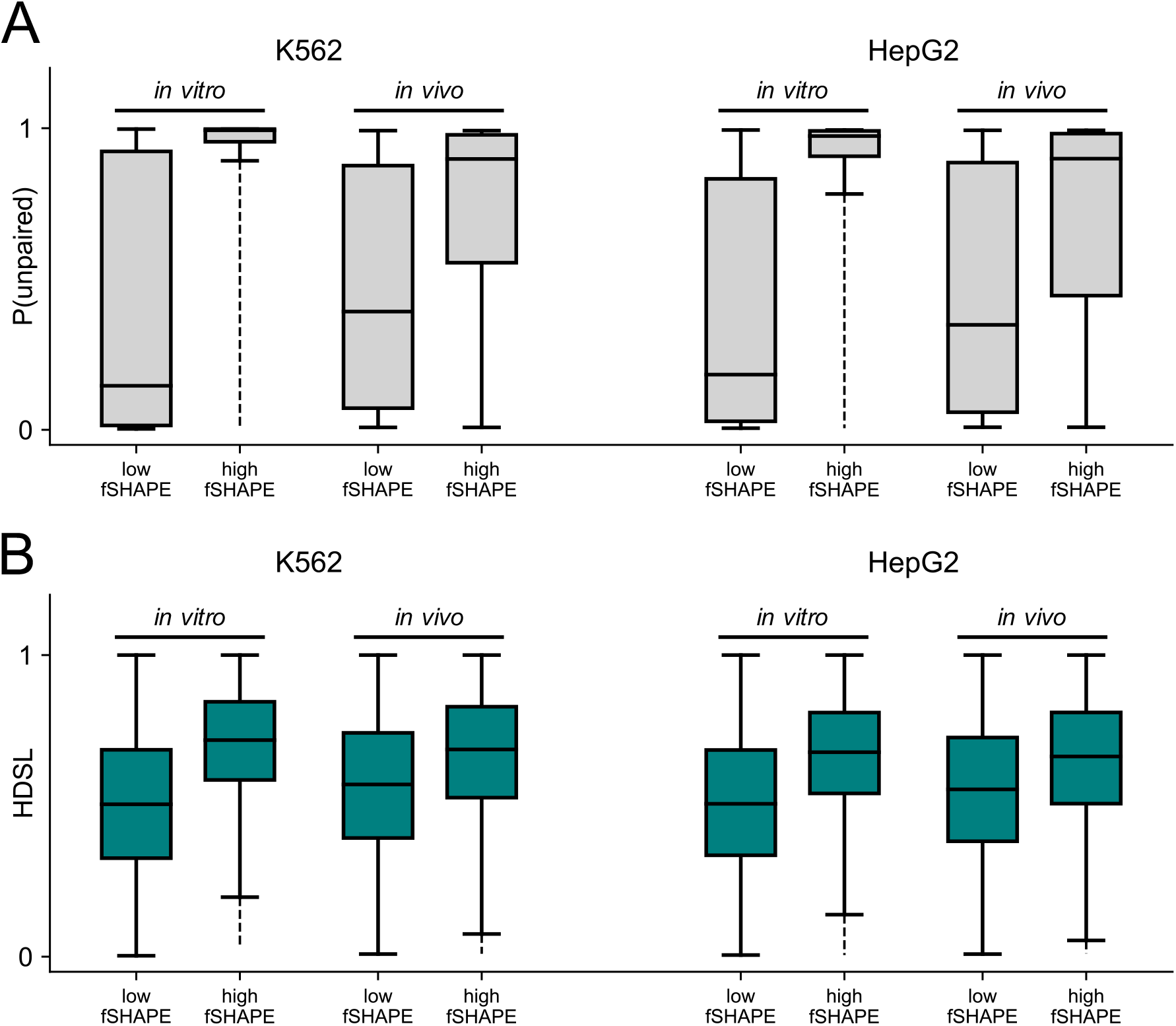
*patteRNA* demonstrates a strong association of RNA structure and RBP binding sites in human cell lines probed as by Corley et al. (32). (A) Unpaired probability boxplots (determined from icSHAPE reactivity via *patteRNA*’s DOM-HMM) for nucleotides with low fSHAPE (fSHAPE < 0) and high fSHAPE (fSHAPE ≥ 2). Within each of the two cell lines, K562 and HepG2, results are presented for both *in vitro* and *in vivo* SHAPE data. (B) HDSL boxplots for nucleotides under the same conditions as (A). Although reactivities indicate that nucleotides likely involved in RBP binding (i.e., nucleotides with high fSHAPE) are remarkedly accessible and therefore likely unpaired, HDSL demonstrates that these reactive nucleotides more frequently occur in the general context of structured regions when compared to nucleotides with low fSHAPE.

However, despite the increased accessibility observed at single nucleotides with high fSHAPE, when one expands the context to the nucleotides’ local neighborhood (i.e., via HDSL analysis), one observes significantly more local structure around nucleotides with high fSHAPE compared to nucleotides with low fSHAPE (**Figure 6B**). This result is consistent with results from NNTM analyses performed by Corley et al., whose interpretation again depended on the computation of Shannon entropy. Our results were achieved without any folding steps and are more statistically significant (*p* < −10^307^for all low/high fSHAPE comparisons in **Figure 6B**, Mann-Whitney *U* test) than originally demonstrated. They were also generated orders of magnitude faster than a comparable NNTM approach, as we will show next. We note that current approaches for summarizing local structuredness from SP data alone, specifically local median reactivity, are generally insufficient for reaching this conclusion (see **Supplementary Figure S7**). This highlights the capability of our method to extract more information from big SP datasets without relying on the additional assumptions and computational overhead of thermodynamic modeling.

### *patteRNA* Processes Large Data Rapidly

An especially appealing property of *patteRNA* is its ability to process big datasets rapidly. To demonstrate its speed in the context of existing methods, we timed our analyses and compared to partition function-based assessment of structure. To this end, we processed the Weeks set, SARS-CoV-2 genome, Mustoe data, and Corley data with three sliding-window partition function analyses of varying computational overhead: partition function calculations with windows of length 3000 nt, spaced 300 nt apart; windows of length 2000 nt, spaced 150 nt apart, and windows of length 150 nt, spaced 15 nt apart. The results of the benchmarks are in Supplementary Figure S8. We observe that *patteRNA* is orders of magnitude faster than sliding-window partition function analysis for massive datasets (e.g., SP data on human transcriptomes). Specifically, *patteRNA* processed the largest dataset included in this study, the Corley data, in less than 1 hour when using a single-threaded implementation (compared to roughly 1 and 7 days for partition function calculation via 150 nt and 2000 nt windows, respectively; 3000 nt window calculations on the Corley data were not performed as they could not be completed in reasonable timeframe). Additionally, our method is natively parallelized, and benchmarks using 12 threads allow *patteRNA* to process such data in less than 10 minutes. Analogous parallelization of partition function-based approaches on large batches of RNA transcripts is relatively simple in theory, but not natively provided “out-of-the-box” for ViennaRNA (meaning it’s up to the user to program their own parallelized calls to the relevant methods). An alternative RNA folding package, RNAstructure (21), does provide scalable parallelization out-of-the-box, but the core folding implementation is about one to two orders of magnitude slower than ViennaRNA. The method was therefore not included in our comparison.

We also compared our method to *RNALfold* (82), an optimized routine within the ViennaRNA package designed to rapidly scan long RNAs for locally-stable structural elements. As expected, we found that this method is capable of processing large data significantly faster than the sliding-window partition function approaches, yet it is nevertheless outpaced by *patteRNA*. Moreover, this method only returns structural elements with sufficiently low free energy (“significantly low” energies judged via an SVM) and, to the best of our knowledge, has not been well-benchmarked against reference structures. Furthermore, *RNALfold* does not attempt to integrate its results to summarize local structuredness, which is key to the type of comparative analyses performed in this study and a central theme of a broad range of recent SP-based studies (32, 47, 71, 72). Nevertheless, this method arrives at a more specific and comprehensive description of local structures (i.e., it can *de-novo* identify stems with bulges and internal loops), whereas *patteRNA*’s analyses here focus specifically on hairpin elements. We note that the incorporation of such local folding routines would likely improve the efficacy of future methods aiming to summarize local structure in large SP datasets, and our results show promising evidence that localized folding can be incorporated without major sacrifices to computational speed.

## DISCUSSION

RNA structure probing experiments are rapidly evolving in terms of their design, scale, and quality. This evolution is accompanied by a need for versatile and scalable methods capable of extracting information from diverse and massive SP data. *patteRNA* is one such tool which was developed to rapidly extract insights from such data. Here, we have demonstrated reformulation of the *patteRNA* framework which increases its speed, adaptability, and precision, enabling it to scale well to data containing millions or billions of nucleotides. Moreover, we have shown that RNA structure can be rapidly quantified and compared in various contexts by detecting the signatures of hairpin elements. Our work expands the repertoire of analyses which *patteRNA* is capable of and demonstrates the power of simpler schemes when interpreting reactivity information. As seen with our benchmarks using a DOM approach, relatively low-resolution discretization schemes (akin to those used to highlight low/medium/high reactivities when visualizing SP data) are valuable when quantifying and mining motifs.

In the context of RNA structure determination, we note that *patteRNA* is not envisioned as a competing method or replacement to traditional NNTM-based approaches. Rather, we view the method as a tool to be used in tandem to RNA folding. As seen in **Figure 3**, NNTM-based methods provide a far more accurate prediction of specific structures and are capable of assessing the entire structure landscape including bulges, internal loops, and internal stems. The analyses via *patteRNA* shown here, on the other hand, intentionally compromise on the type of structures considered in the analysis in order to maximize the speed and scalability of the approach. As a consequence, *patteRNA* is most useful when assessing large-scale data. For instance, as we demonstrated, it could be utilized to quantify macroscopic structural trends related to specific regions, or it could be used to identify regions of RNA which see differential structuredness associated to some factor, which might then be followed by more intensive RNA folding approaches (e.g., partition function computation). In this way, *patteRNA* helps mitigate the computational limitations of such methods, especially for those who do not have advanced computing hardware at their disposal. Finally, although analyses in this study generally focus on using *patteRNA* to derive information on structuredness via hairpins, the method itself is fundamentally a versatile structure-mining algorithm which has been demonstrated to effectively search for putative functional motifs across in transcriptome-wide data (57).

Our analysis of the SARS-CoV-2 5’UTR is distinguished from the others by a comparison of HDSL with specific structures that have been validated in a plethora of ways, including NMR spectroscopy (78). We remarked on a great correspondence of HDSL peaks and stable structural elements, indicating that HDSL captures more than just local structure—it retains information on specific motifs with high resolution. This observation is important in the context of our analysis of Corley et al.’s fSHAPE data. Namely, the increase in HDSL around sites with high fSHAPE (**Figure 6B**) suggests the possibility that RBP frequently associate not only in the context of structured regions, but specifically in the context of hairpin-like elements. RBP which recognize sequence motifs in hairpin-loops have previously been identified (83, 84), but our results demonstrate the plausibility that the association between hairpin elements and RBP is more prevalent than previously thought. This is not entirely unexpected, as RBP are known to bind both dsRNA and ssRNA in a manner that correlates with the structure of the protein (85). Moreover, RBP binding ssRNA are observed to associate at unpaired bases stemming from RNA helix irregularities (e.g., bulges and internal loops) (86), also placing them in the context of hairpin elements. Recent studies have further documented that structured RNAs interact with a larger number of proteins than less structured RNAs (85). Our result further strengthens the utility of *patteRNA* in mining biologically relevant structures transcriptome-wide.

Looking ahead to future development of rapid analysis of SP data, *patteRNA* is well-suited to adapt to evolving probing technologies and datasets. That being said, its current implementation does come with several limitations. First, motif mining depends on the definition of specific secondary structures, which limits its application to situations where a specific structure or small collection of similar structures can be defined. For motifs like hairpins, this means that considering situations where a bulge or internal loop may or may not be present complicates analyses due to the combinatorial explosion of unique secondary structures needed to define all possible hairpin architectures through loop size, bulge size, and bulge position. *patteRNA* is already capable of exhaustively mining such motifs, but such analyses come at the cost of significant computational overhead, generally working against the utility of the method. A more efficient approach for motif mining which naturally considers alternative similar structures within a region could theoretically address some parts of this limitation. Secondly, although the circumvention of RNA folding enables rapid computational analyses, it also handicaps the accuracy of the approach, as the energetic favorability of sequences within stems and loops is ignored. The incorporation of an optimized local folding routine could likely assist in this regard, although the coupling of such models into a statistical model like *patteRNA* is non-trivial. Nevertheless, methods like *RNALfold* (82) bode for the potential incorporation of NNTM-derived information without sacrificing on speed and scalability. Regardless of these limitations, however, *patteRNA* remains a viable computational method for the rapid assessment and quantification of structural trends in the largest SP datasets.

## DATA AVAILABILITY

The latest version of *patteRNA*, version 2.0, was used for all analyses in this study. *patteRNA* is an open-source Python 3 module and is freely available at www.github.com/AviranLab/patteRNA under the BSD-2 license. Python scripts for generating simulated datasets, computing statistical benchmarks (e.g., ROC and PRC), and post-processing of HDSL profiles related to genes in the Mustoe data are available in **Supplementary File 1**. The original datasets used in this study are all publicly available from the indicated references in **Table 1**.

## SUPPLEMENTARY DATA

Supplementary Data are available online.

- Supplementary Figures and Tables (.pdf)
- Supplementary Material (.pdf)
- Supplementary File 1 (.zip)

